# Taxometer: Improving taxonomic classification of metagenomics contigs

**DOI:** 10.1101/2023.11.23.568413

**Authors:** Svetlana Kutuzova, Mads Nielsen, Pau Piera, Jakob Nybo Nissen, Simon Rasmussen

## Abstract

For taxonomy based classification of metagenomics assembled contigs, current methods use sequence similarity to identify their most likely taxonomy. However, in the related field of metagenomics binning contigs are routinely clustered using information from both the contig sequences and their abundance. We introduce Taxometer, a neural network based method that improves the annotations and estimates the quality of any taxonomic classifier by combining contig abundance profiles and tetra-nucleotide frequencies. When applied to five short-read CAMI2 datasets, it increased the average share of correct species-level contig annotations of the MMSeqs2 tool from 66.6% to 86.2% and reduced the share of wrong species-level annotations in the CAMI2 Rhizosphere dataset two-fold on average for Metabuli, Centrifuge, and Kraken2. Finally, we applied Taxometer to two complex long-read metagenomics data sets for benchmarking taxonomic classifiers. Taxometer is available as open-source software and can enhance any taxonomic annotation of metagenomic contigs.

## Main

Metagenomic classifiers annotate reads or contigs with taxonomic information by searching for similar substrings in a collection of reference sequences. The quality of annotations depends on the chosen method ^1–8^ as well as the reference database, requiring a careful selection of tools depending on the research context. Due to the high complexity of metagenomics data and inevitable database incompleteness, full sample characterization is not usually achieved.

Most metagenomic classifiers search for each string individually without taking advantage of the dataset context^9^. However, for the related problem of metagenomics binning, state-of-the-art methods rely on abundances and tetra-nucleotide frequencies (TNFs) to link contigs of the same origin ^10–15^. Similarities in the feature vectors across contigs in the dataset allows reference-free binning of the contigs to reconstruct metagenome-assembled genomes (MAGs) from poorly studied environments with insufficient database representation ^16–18^.

We propose a method to enhance taxonomic profiling by utilizing the features that are used for binning. Our method, Taxometer, is designed to run on any standard metagenomics dataset of one or more samples where annotation of the contigs is required. Taxometer is based on a neural network that uses TNFs, abundances, and taxonomic labels from any metagenomic classifier. The neural network is then trained to predict taxonomy for this particular dataset using the subset of contigs that have an annotation. Finally, the trained network is applied to the input contigs resulting in refined taxonomic labels of contigs as well as annotation of contigs without taxonomic labels (Figure 1a). As the taxonomic labels contain the full path from the taxonomic root to the placement on the bacterial phylogenetic tree, we implemented a tree based hierarchical loss. This allows partial annotations not covering all taxonomic levels and that Taxometer can predict across all taxonomic levels accompanied by scores ranging from 0.5 to 1. The workflow results in refined taxonomy annotations, containing more true positives and fewer false positives compared to the input taxonomy classification. Importantly, as contig abundances are specific to a certain dataset we train and apply Taxometer to each dataset. Therefore, even though Taxometer is designed as a supervised deep learning method we do not transfer the model or try to generalize across datasets.

**Fig. 1.**
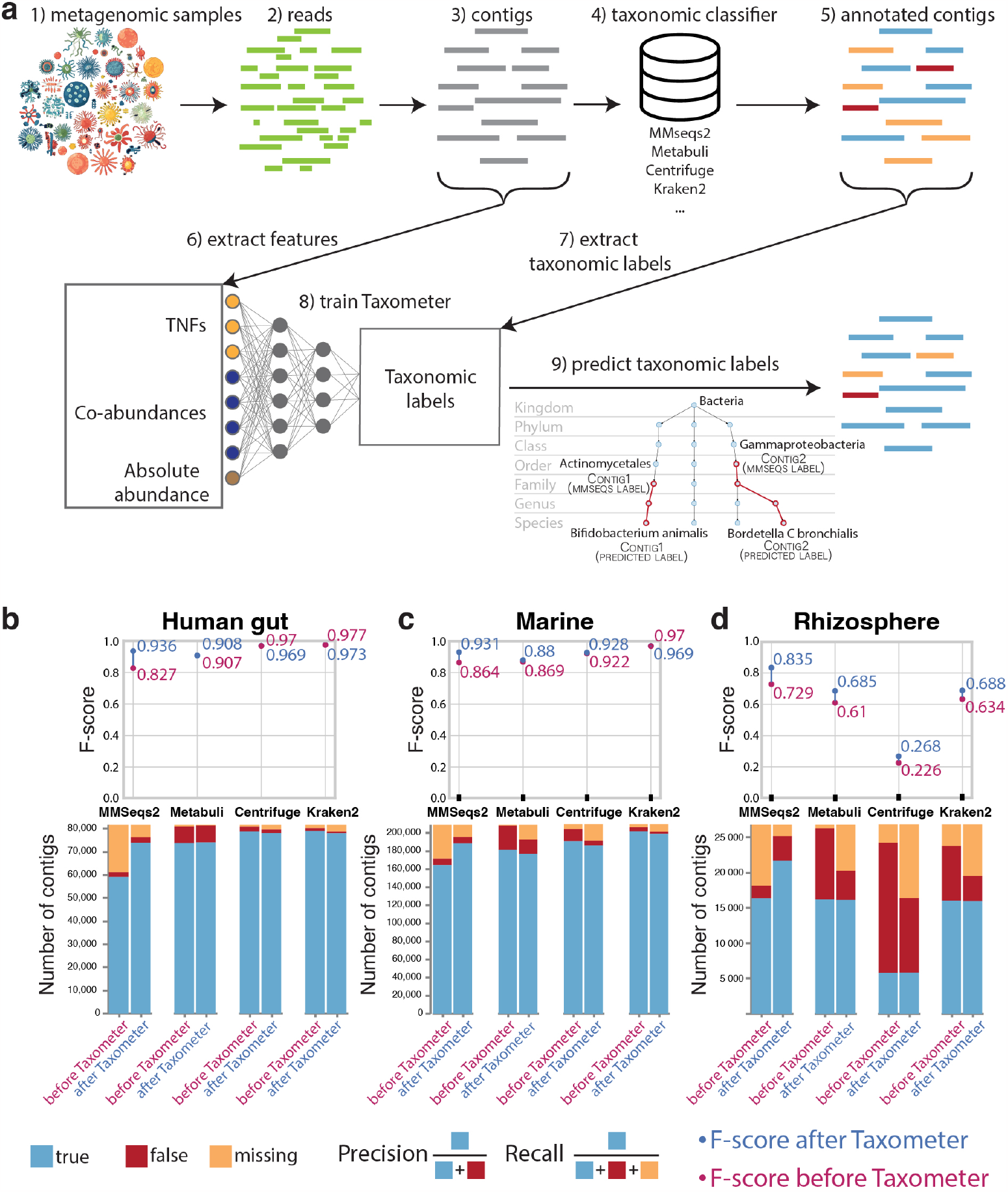
Workflow and CAMI2 results. **a**, Taxonomic profiling workflow using Taxometer. After assembling the metagenomic reads, contigs are annotated with any taxonomic classifier. Contigs are processed via Taxometer to extract abundances and tetra-nucleotide frequencies. Taxometer is then trained on the extracted features to predict the taxonomic annotations. A hierarchical loss allows training the model on the annotations from all levels. The predictions are then made for all taxonomic levels for each contig (e.g. completing the annotation from Actinomycetales order to Bifidobacteruim animalis species). For each label, a score is provided. **b**,**c**,**d** Taxonomic classifier annotations and Taxometer results at species level, compared to the ground truth, score threshold 0.95, CAMI2 Gastrointestinal, Marine and Rhizosphere datasets. MMseqs2 and Metabuli returned GTDB annotations, Centrifuge and Kraken2 returned NCBI annotations. The comparisons are to the ground truth labels.

To demonstrate that Taxometer could improve the annotations of different taxonomic classifiers, we trained it on MMseqs2^3^ and Metabuli ^19^ configured to use the GTDB database ^20^ and on Centrifuge ^21^ and Kraken2^22^ configured to use the NCBI database ^23^. For the CAMI2 human microbiome datasets we found that MMseqs2 correctly annotated on average 66.6% of contigs at species level (Figure 1b). When applying Taxometer, trained on the MMseqs2 annotations, the amount of correct annotated contigs increased to 86.2%. Additionally, when applied to two more challenging datasets, CAMI2 Marine and Rhizopshere, Taxometer increased the MMseqs2 annotation level from 78.6% to 90%, and from 61.1% to 80.9%, respectively (Figure 1b). This was reflected in F1-scores of the annotation improvement between 0.1 and 0.13 for the human microbiome datasets. When we applied Metabuli, Centrifuge, and Kraken2 to the CAMI2 human microbiome datasets, they correctly annotated on average more than 94.8% of contigs at species level. Here, Taxometer did not improve on this close-to-perfect annotation level but also did not decrease performance (F1-score absolute change *<*0.002) (Supplementary Data 1). However, when applied to the Marine and Rhizosphere datasets, the performance of the three methods was much lower. For instance, Metabuli provided wrong species annotations for 12.7% and 37.6% of the two datasets, respectively (Figure 1d, Supplementary Figure S1). Here, Taxometer reduced the number of wrong annotations to 7.6% and 15.4%, increasing F1-score from 0.87 to 0.88 and from 0.61 to 0.69 respectively. Similarly, for Centrifuge and Kraken2 applied to the Rhizosphere dataset, Taxometer reduced the amount of wrong annotations from 68.7% to 39.5% (F1-score from 0.22 to 0.27) and from 28.7% to 13.3% (F1-score from 0.64 to 0.68), respectively (Supplementary Figure S1, Supplementary Data 1). Even though Taxometer had high precision for taxonomic annotations it made mistakes and re-annotated a small amount of the correct MMseqs2 species-level annotations incorrectly (1.6%-3.3% on the CAMI2 human microbiome). Nonetheless, Taxometer can fill annotation gaps and remove incorrect taxonomic labels of large numbers of contigs from diverse environments, while only mislabeling a small minority of correctly labeled contigs.

Given the ability of Taxometer to correctly predict new as well as correct wrong annotations we investigated the contribution of abundance and TNFs features for the predictions. We, therefore, trained Taxometer to predict MMseqs2 annotations for the CAMI2 human microbiome datasets using either the abundances or TNFs (Figure 2a, Supplementary Figure S2). Here we found that for higher taxonomic levels (phylum to genus) up to 98% of Taxometer annotations could be reproduced by training using only TNFs. This is in concordance with previous findings that TNFs could be used to classify metagenomics fragments at the genus level and that abundance showed better strain-level binning performance compared to TNFs ^15,24^. The number of correct species labels predicted by the model that combined both TNFs and abundances was 18%-35% larger than the models that only used TNFs or abundances for MMseqs2 annotations of the CAMI2 Airways dataset. Additionally, we investigated the importance of varying the threshold of the annotation score. Here, we found that a threshold score of 0.95 provided a good balance between recall and precision (Supplementary Figure S3, Supplementary Figure S4).

**Fig. 2.**
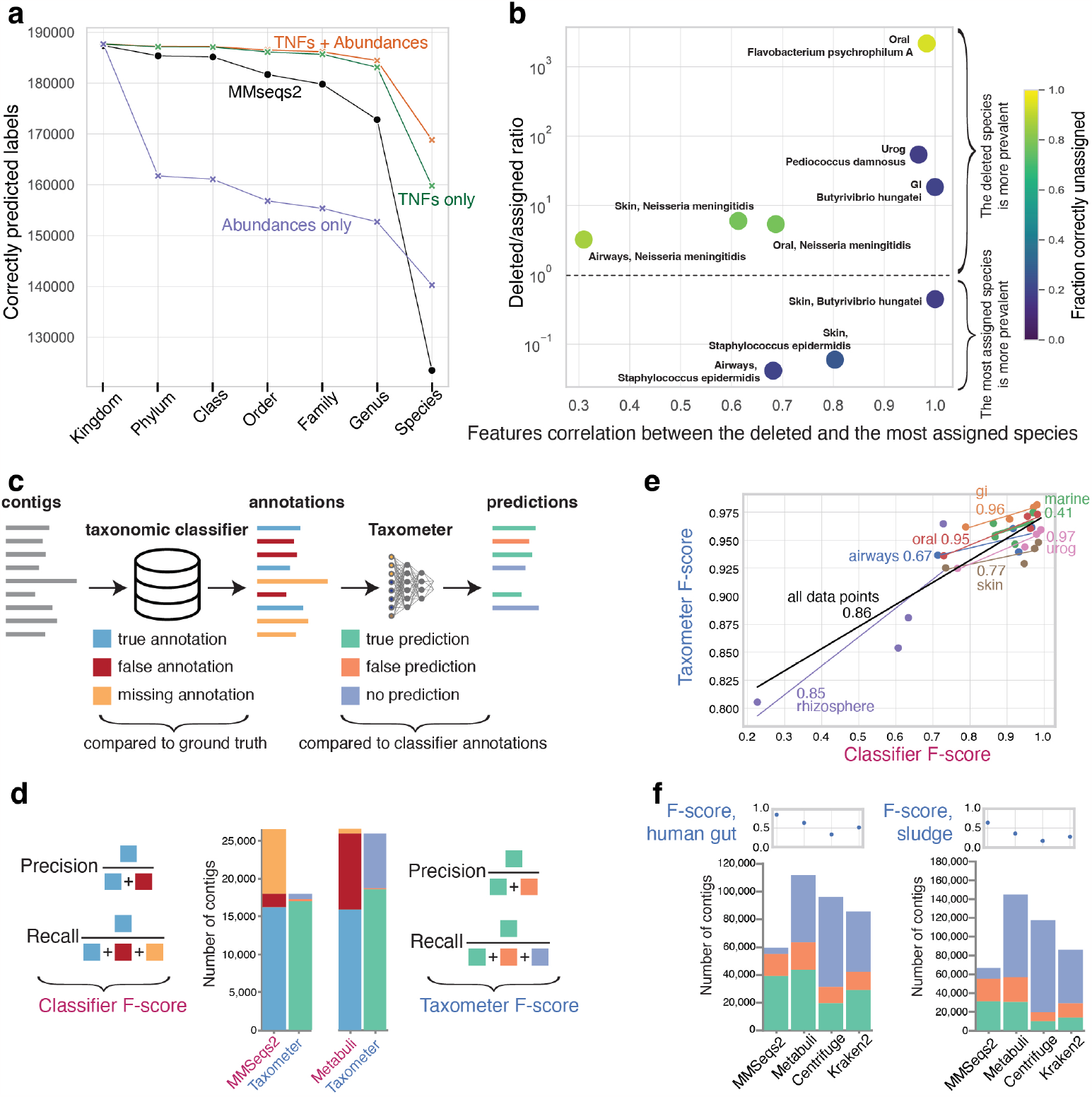
Analysis of feature importance, novel taxa and long-read datasets. **a**, Contribution of abundances and TNFs features to Taxometer performance demonstrated on the CAMI2 Airways short-read dataset. The amount of correctly predicted contigs labels at each taxonomic level using a score threshold of 0.5. **b**, Simulation analysis of unknown taxa. X-axis: Pearson correlation coefficient between the mean feature vectors of the deleted and the assigned species. Y-axis: ratio between the number of contigs of the deleted species (“deleted”) and the number of contigs of the species that was the most prevalent among the incorrectly assigned (“assigned”) in the training set. The color legend shows the share of correctly missing labels, equal to 1 − *F P*, where FP is the share of false positives. FP is high when the assigned species was more prevalent in the training set and TNFs and abundances are highly correlated between the deleted and the assigned species. **c**, K-fold evaluation description. The Taxometer predictions are compared to the classifer annotations, not the ground truth labels. **d**, An example of the number of true positives, false positives, and false negatives used in the k-fold evaluation, species level. The total number of contigs for Taxometer predictions equals the number of annotations initially returned by a classifier. **e**, F1-scores of Taxometer and classifiers for each dataset. X-axis is the F1-score of a taxonomic classifier compared to ground truth, Y-axis is the F1-score of Taxometer predicting the taxonomic classifier annotations, score threshold 0.5, all data points.**f**, Real long-read datasets k-fold evaluation for human gut and sludge environments.

To explore the limitations of Taxometer we investigated the performance when species in the dataset were missing from the database. To achieve this we deleted annotations from five species in the CAMI2 human microbiome MMseqs2 results before training Taxometer. This resulted in removing between 649 and 5,127 contig annotations per dataset. As the deleted species were not in the training set, a perfect classifier should assign missing labels to the contigs that belong to this species. In our experiments, Taxometer predicted the correct genus label for all these contigs. However, the share of incorrectly assigned annotations at species level varied between 6% and 82% across the species and the datasets. For these wrong annotations, we found that the number of contigs of the deleted and the assigned species, and the mean feature correlation between them were the most important factors (Figure 2b). We found that false positives tended to occur when the assigned species were more prevalent in the training set than the deleted species, e.g. in the Airways dataset *Staphylococcus aureus* was assigned to 1,312 of 1,879 contigs from the deleted species *Staphylococcus epidermidis*. That can be explained by *Staphylococcus epidermidis* being 17 times more prevalent in the training set than *Staphylococcus aureus*, with 78% of all contigs from the *Staphylococcus* genus were from *Staphylococcus aureus*. Second, we found it to make mistakes when TNFs and abundances were highly correlated between the deleted and the assigned species. For instance, in the Gastrointestinal dataset, 183 out of 229 contigs of the deleted *Butyrivibrio hungatei* species were assigned to *Butyrivibrio sp900103635*, that only had 8 contigs in total. However, the pearson correlation of the mean feature vectors for these two species was 0.99. Thus, for the contigs of a novel taxon, Taxometer might assign a closely related taxon instead of returning a missing annotation.

We were interested in evaluating the performance of Taxometer for the annotation of long-read based metagenomics. However, benchmark datasets of sufficient size and complexity were not available. We, therefore, investigated if the consistency of Taxometer annotations could be used as a measure of classifier performance when ground truth labels were missing. Because metagenomics binning has been able to generate hundreds of thousands of MAGs, TNFs and abundances carry a strong signal for contigs of the same origin ^18^. Thus, we hypothesized that the ability of Taxometer to predict classifier annotations could reflect the performance of the classifier. Specifically, the more incorrect labels a classifier assigns to contigs of the same origin, the harder it will be for Taxometer to reproduce the labels assigned by a classifier. To investigate this we acquired annotations of four classifiers for the seven CAMI2 datasets, resulting in 28 sets of taxonomic labels. We divided each dataset into five folds and trained Taxometer five times using a new fold as the validation set. We then compared Taxometer predictions to classifier annotations and classifier annotations to the ground truth (Figure 2c,d). We found that Taxometer and classifier F1-score metrics were correlated with Pearson Correlation Coefficient (PCC) of 0.86 across all datasets (Figure 2e, Supplementary Figure S5). Thus, in the absence of ground truth labels the F1-score of Taxometer prediction for classifier labels can be used as a benchmark for different taxonomic classifiers within a dataset.

We then identified two PacBio HiFi read datasets of complex metagenomes; 4 samples from the human gut microbiome and 3 samples from an anaerobic digestion reactor sludge ^25^. These datasets did not have ground truth labels and when we applied either GTDB- or NCBI-based classifiers they disagreed for 28%-39% of the contigs at species level annotations (Supplementary Figure S6). Therefore, we applied the k-fold evaluation scheme described above to use Taxometer F1-scores for benchmarking the classifiers. Here we found that MMseqs2 had the highest F1-score, i.e. 0.85 for the human gut dataset (Metabuli 0.64, Centrifuge 0.34, Kraken2 0.52) (Figure 2f, Supplementary Figure S7a). The range of precision and recall were similar as when evaluated on the CAMI2 data for the contigs with missing MMseqs2 species annotations, approximately [0.8, 0.95] for precision and [0.6, 0.85] for recall (Supplementary Figure S7b). This was consistent with the performance of the classifiers on the most difficult CAMI2 dataset, Rhizosphere, and thus we concluded that MMseqs2 returned the most correct annotations for both the human gut microbiome and sludge datasets compared to any of the other classifiers.

Finally, Taxometer is designed as a lightweight tool that is less compute-intensive than the taxonomic annotations themselves. For instance, annotation of the CAMI2 and long-read datasets with MMSeqs2 took 2-4 hours, while training Taxometer with a single GPU took less than 30 minutes for all datasets (Supplementary Figure S8, Supplementary Data 1). Similarly, training could be done using CPUs in 10-120 minutes.

In summary, Taxometer can improve taxonomic annotations of any contig-level metagenomic classifier. Tested on the annotations of the two taxonomic classifiers across several real and simulated datasets, Taxometer both filled annotation gaps and deleted incorrect labels. Additionally, Taxometer provides a metric for evaluating the quality of annotations in the absence of ground truth. The analysis of the long-read datasets suggested that the performance was stable across sequencing technologies. The main limitation of applying Taxometer to a metagenomics dataset is that read-level data is needed to calculate contig abundances for a particular dataset. Taxometer therefore cannot be used to annotate contigs downloaded from a large database without such information. However, taxonomic annotations can be improved in any metagenomics experiment where reads are available. This could be a metagenomics experiment run in-house in a lab or where corresponding read-level data is available for download from SRA or ENA. Therefore, we believe that Taxometer is broadly applicable for improving annotations across almost all metagenomics datasets.

## Methods

### Evaluation datasets

For the short-read benchmarks, we used synthetic CAMI2 datasets from five human microbiomes (Airways, Oral, Skin, Urogenital, and Gastrointestinal) and two environmental microbiomes (Marine and Rhizosphere) ^26^. For the CAMI2 evaluations, we compared the results of taxonomic classifiers and Taxometer to the provided ground truth labels for each contig. We also benchmarked Taxometer using two real PacBio HiFi long-read datasets, a ‘human gut’ dataset from human stool samples ^25^ and a ‘sludge’ dataset from anaerobic digestion reactor sludge (ENA accessions ERR10905741-ERR10905743). In the absence of a synthetic long-read dataset with ground truth labels, the evaluations were done on real datasets via k-fold evaluation.

### Data preprocessing

For short-read benchmarking, we used sample-specific assemblies for the seven CAMI2 datasets: Airways (10 samples), Oral (10 samples), Skin (10 samples), Gastrointestinal (10 samples), Urogenital (9 samples), Marine (10 samples), Rhizosphere (21 samples). For each dataset we aligned the synthetic short paired-end reads from each sample using bwa-mem (v.0.7.15) ^27^ to the concatenation of per-sample contigs from the particular dataset. BAM files were sorted using samtools (v.1.14) ^28^. For all datasets, we used only contigs *>*2,000 base pairs (bp). For long-read benchmarking we used a human gut microbiome dataset with 4 samples and a dataset from anaerobic digester sludge with 3 samples ^29^, both sequenced using Pacific Biosciences HiFi technology. We assembled each sample using metaMDBG (v. b55df39) ^30^, mapped reads using minimap2 2.24^31^ with the ‘-ax map-hifi’ setting, and from there proceeded as with the short reads.

### Abundances and TNFs

Computation of abundances and TNFs was done using the VAMB metagenome binning tool ^15^. To determine TNFs, tetramer frequencies of non-ambiguous bases were calculated for each contig, projected into a 103-dimensional orthonormal space and normalized by z-scaling each tetranucleotide across the contigs. To determine the abundances of each sample, we used pycoverm 0.6.0^32^. The abundances were first normalized within sample by total number of mapped reads, then across samples to sum to 1. To determine absolute abundance, the sum of abundances for a contig was taken before the normalization across samples. The dimensionality of the feature table was then *N*_*c*_ *×* (103 + *N*_*s*_ + 1) where *N*_*c*_ was the number of contigs, *N*_*s*_ was the number of samples.

### CAMI2 ground truth labels

For the five CAMI2 human microbiome datasets, the provided genomes sequences were classified with GTDB-tk ^33^ tool which resulted in all the contigs annotated to species level with GTDB ^20^ identifiers. Ground truth labels for Marine and Rhizosphere datasets were converted from NCBI ^34^ to GTDB v207 using the *gtdb to taxdump* tool (v. 24b82d6) available at https://github.com/nick-youngblut/gtdb_to_taxdump. Not all NCBI labels had a 1-to-1 match in GTDB and not all contigs were annotated in the CAMI2 dataset to the species level. The resulting number of fully annotated contigs used in the analysis was 208,783 out of 438,686 contigs (48%) for the Marine dataset and 26,734 out of 300,222 contigs (9%) for the Rhizosphere dataset.

### Taxonomic classifiers

We obtained the taxonomic annotations for contigs of all seven short-read and two long-read datasets from MMseqs2 (v.7e2840) ^3^, Metabuli (v.1.0.1) ^19^, Centrifuge (v1.0.4) ^21^ and Kraken2 (v2.1.3) ^22^. For MMseqs2, we used the *mmseqs taxonomy* command. For Metabuli, we used the *metabuli classify* command with *–seq-mode 1* flag. For Centrifuge, we used the *centrifuge* command with *-k 1* flag. For Kraken2, we used the *kraken* command with *–minimum-hit-groups 3* flag. MMseqs2 and Metabuli were configured to use GTDB v207 as the reference database. Centrifuge and Kraken2 were configured to use NCBI identifiers.

### Network architecture and hyperparameters

The input vector of dimensionality *N*_*c*_ *×* (103 + *N*_*s*_ + 1) was passed through 4 fully connected layers ((103+*N*_*s*_ +1)*×*512, 512*×*512, 512*×*512, 512*×*512) with leaky ReLU activation function (negative slope 0.01), each using batch normalization (epsilon 1*e*− 05, momentum 0.1) and dropout (*P* = 0.2). The output layer had dimensionality 512*× N*_*l*_ where *N*_*l*_ was the number of leaves in the taxonomic tree (see subsection 4). For all datasets, the network was trained for 100 epochs with batch size 1024 using the Adam optimizer with learning rates set via D-Adaptation ^35^. The model was implemented using PyTorch (v.1.13.1) ^36^, and CUDA (v.11.7.99) was used when running on a V100 GPU.

### Hierarchical loss

A phylogenetic tree was constructed for each dataset from the taxonomy classifier annotations for the set of contigs. Thus, the resulting taxonomy tree *T* was a subgraph of a full taxonomy and the space of possible predictions was restricted to the taxonomic identities that appeared in the search results. For the above experiments, we used a flat softmax loss. Let *N*_*l*_ be the number of leaves in the tree *T*. The likelihoods of leaf nodes of the taxonomy tree were obtained from the softmax over the network output layer with dimensionality 1 *× N*_*l*_. The likelihood of an internal node was then a sum of likelihoods of its children and computed recursively bottom-up. The model output was a vector of likelihoods for each possible label. For the backpropagation the negative log-likelihood loss was computed for all the ancestors of the true node and the true node itself. Predictions were made for all taxonomic levels and for each level, the node descendant with the highest likelihood was selected. If no node descendant had likelihood *>* 0.5, the predictions from this level and the levels below were not included in the output.

### K-fold evaluation

All the contigs in a dataset that were annotated by a classifier are randomly divided into five folds. Taxometer was then trained five times, each time using one fold as a validation set and the remaining four folds as training set. The predictions were made for the five validation sets after training the network on the remaining four. This way the prediction for each contig was made without the prior knowledge of the classifier annotation of this contig. The F1-score metric that was used to benchmark the classifiers within a dataset was then calculated as following:

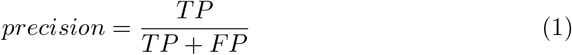

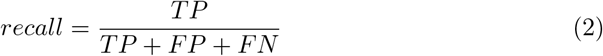

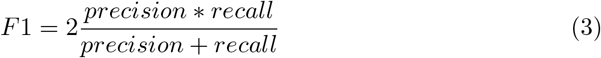

where *TP* (true positives) is the total number of contigs for which the Taxometer prediction is the same as the classifier annotation, *FP* (false positives) is the total number of contigs for which the Taxometer prediction is different from the classifier annotation and *FN* (false negatives) is when Taxometer prediction is missing, but the classifier annotation exists. For this evaluation, a score threshold of 0.5 was used.

## Acknowledgements

S.K., M.N. and S.R. were supported by the Novo Nordisk Foundation (grant NNF19SA0059348). P.P, J.N.N and S.R. were supported by the Novo Nordisk Foundation (grant NNF20OC0062223). S.K., P.P, J.N.N and S.R. were supported by the Novo Nordisk Foundation (grant NNF14CC0001). S.R. was supported by the Novo Nordisk Foundation (grant NNF21SA0072102).

## Author Contributions

S.K, J.N.N and S.R designed the experiments. P.P and J.N.N preprocessed the datasets. S.K. wrote the software and performed the analysis. M.N, P.P, J.N.N and S.R. provided guidance and input for the analysis. S.K. wrote the manuscript with contributions from all coauthors. All authors read and approved the final version of the manuscript.

## Data availability

The CAMI2 datasets were downloaded from https://data.cami-challenge.org/participate from “2nd CAMI Toy Human Microbiome Project Dataset” (5 human microbiome datasets), “2nd CAMI Challenge Marine Dataset” (Marine), “2nd CAMI Challenge Rhizosphere challenge” (Rhizosphere). The long-read human gut dataset is available at https://downloads.pacbcloud.com/public/dataset/Sequel-IIe-202104/metagenomics/. The long-read sludge dataset is available at the ENA as part of the study PRJEB39861.

## Code availability

All code can be found on GitHub at https://github.com/RasmussenLab/vamb and is freely available under the permissive MIT license.

