## Supplementary Figures for "Taxometer: Improving taxonomic classification of metagenomics contigs"

### List of Figures

S1 Rhizosphere annotation and prediction quality for all domains . . . . . S2

S2 Contribution of abundances and TNFs features to Taxometer performance S3

S3 Precision-recall curves for CAMI2 human microbiome . . . . . . . . . S4

S4 Precision-recall curves for CAMI2 marine and rhizosphere . . . . . . . S5

S5 F-scores of Taxometer and classifiers for each dataset . . . . . . . . . . S6

S6 Long-read datasets classification discrepancies . . . . . . . . . . . . . . S7

S7 K-fold results for the long-read datasets . . . . . . . . . . . . . . . . . S8

S8 GPU runtimes for all datasets . . . . . . . . . . . . . . . . . . . . . . . S9

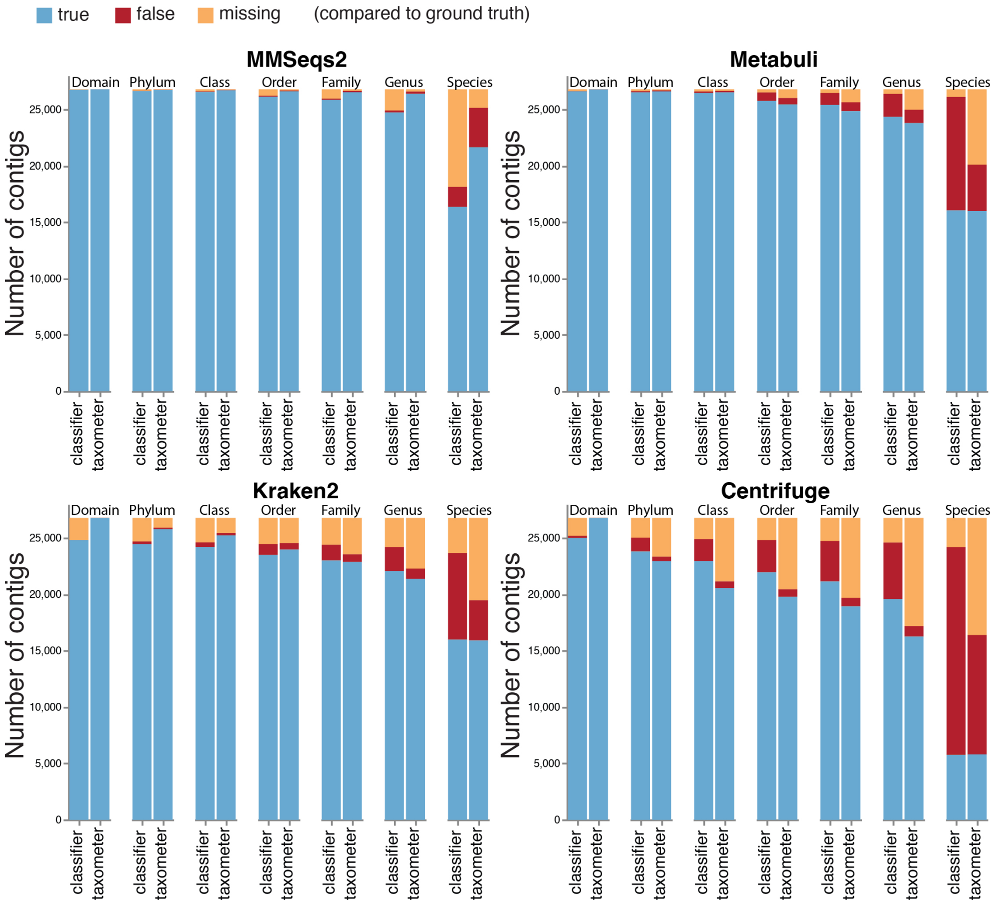


**Supplementary Figure S1 Rhizosphere annotation quality and prediction quality for all domains.** The number of true, false and missing annotations for four taxonomic classifiers and predictions of Taxometer trained on each classifier, compared to ground truth.

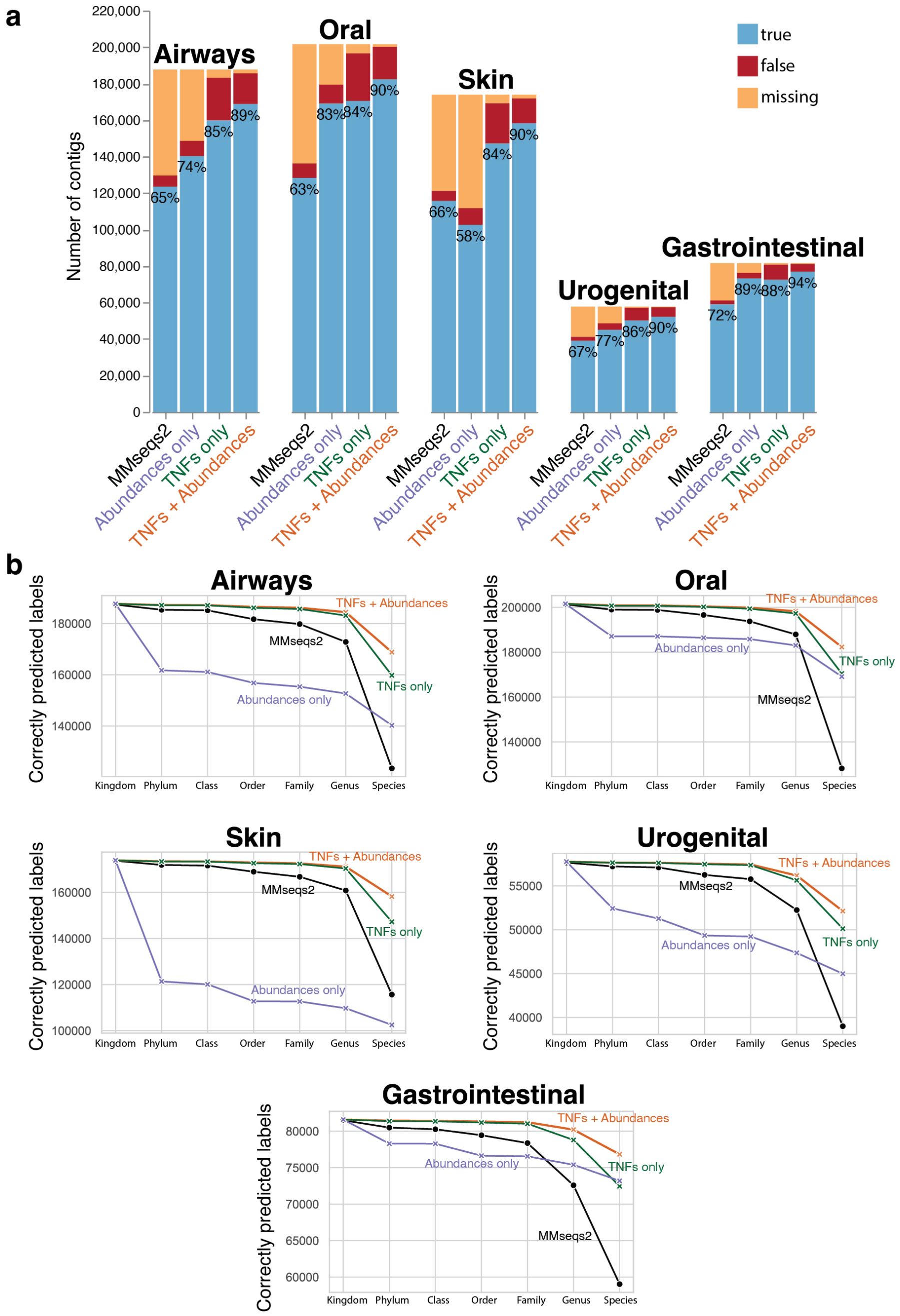


**Supplementary Figure S2 Contribution of abundances and TNFs features to Taxometer performance. a,** Species-level predictions compared to ground truth with score threshold 0.5, CAMI2 human microbiome short-read dataset. **b,** Number of correctly predicted labels on each taxonomic level.

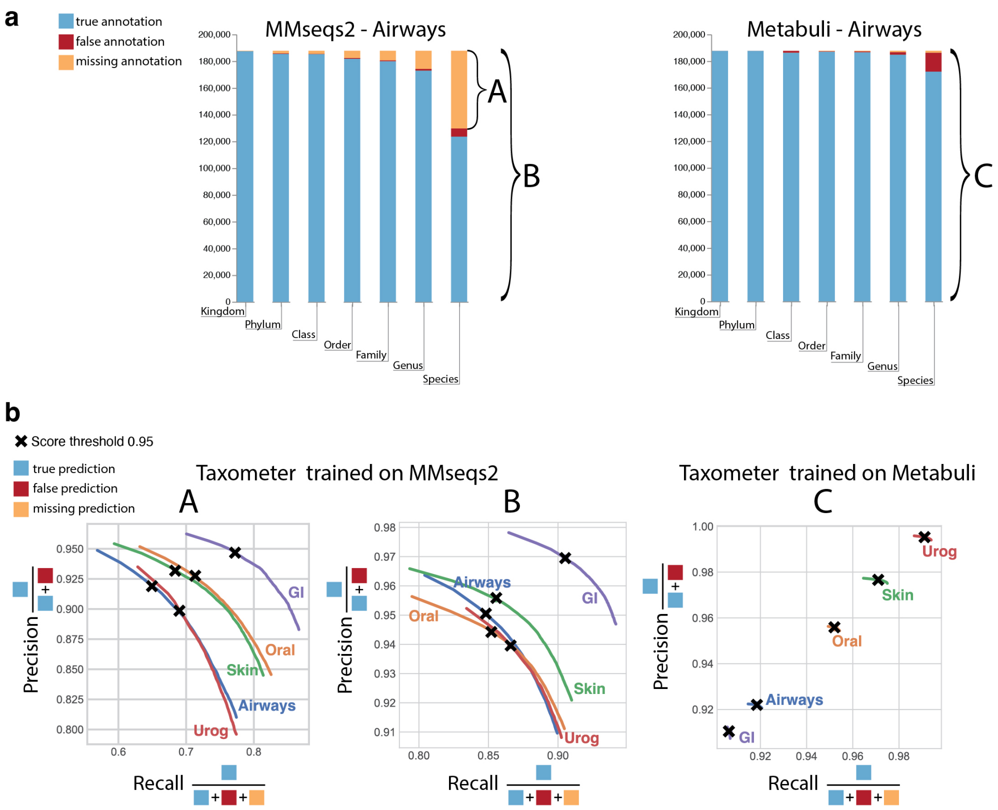


**Supplementary Figure S3 Precision-recall curves for CAMI2 human microbiome. a,** The number of true, false and missing MMseqs2 and Metabuli annotations for CAMI2 Airways dataset. **b,** Precision-recall curves for Taxometer scores in the range $[0.5, 1]$ for species labels calculated for A-labelled contigs with missing species-level MMseqs2 annotations; B- and C-labelled was based on all contigs. For A and B, Taxometer was trained on MMseqs2 annotations, and for C using Metabuli annotations. The cross marks the score threshold 0.95.

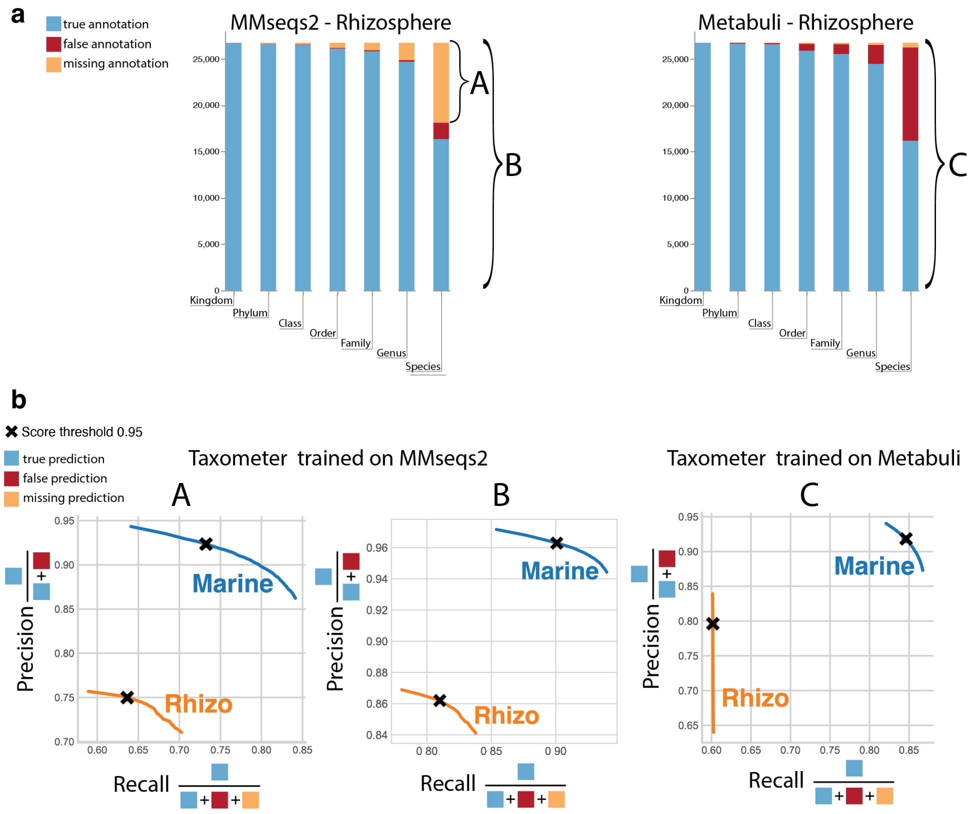


**Supplementary Figure S4 Precision-recall curves for CAMI2 marine and rhizosphere.** **a,** The number of true, false and missing MMseqs2 and Metabuli annotations for CAMI2 Rhizosphere dataset. **b,** Precision-recall curves for Taxometer scores in the range [0.5, 1] for species labels calculated for A-labelled contigs with missing species-level MMseqs2 annotations; B- and C- labelled was based on all contigs. For A and B, Taxometer is trained on MMseqs2 annotations, for C, on Metabuli annotations. The cross marks the score threshold 0.95.

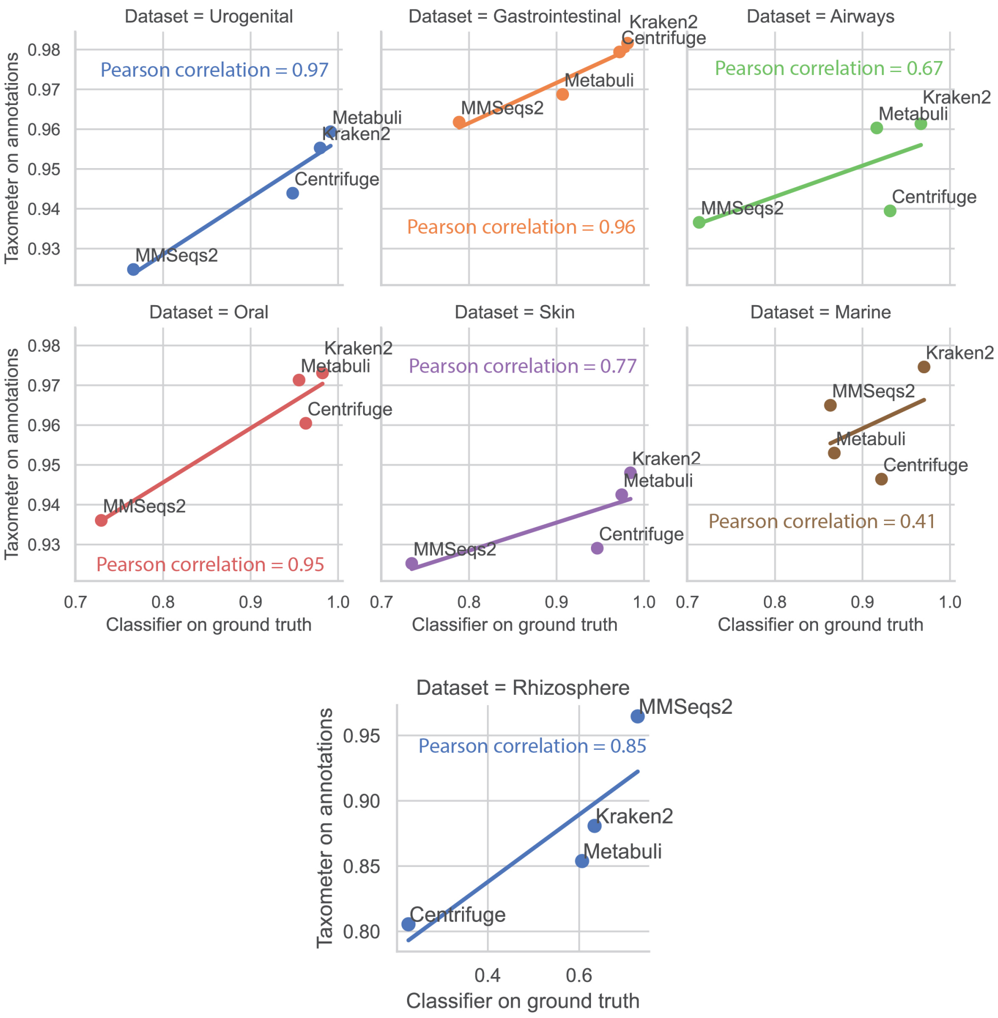


**Supplementary Figure S5 F-scores of Taxometer and classifiers for each dataset.** X-axis is F-score of a taxonomic classifier compared to ground truth, Y-axis is F-score of Taxometer predicting the taxonomic classifier annotations, score threshold 0.5, on a subplot per dataset.

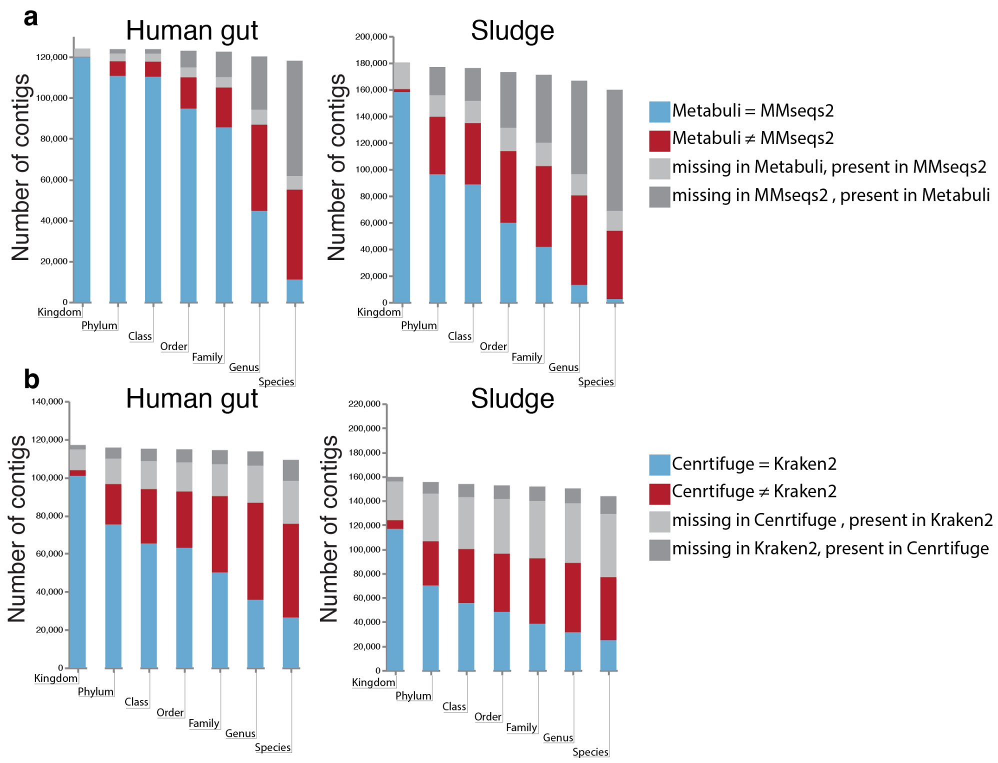


**Supplementary Figure S6 Long-read datasets classification discrepancies. a,** Discrepancies between long-read contigs annotations of GTDB classifiers, MMseqs2 and Metabuli, on all taxonomic levels. **b,** Discrepancies between long-read contigs annotations of NCBI classifiers, Kraken2 and Centrifuge, on all taxonomic levels.

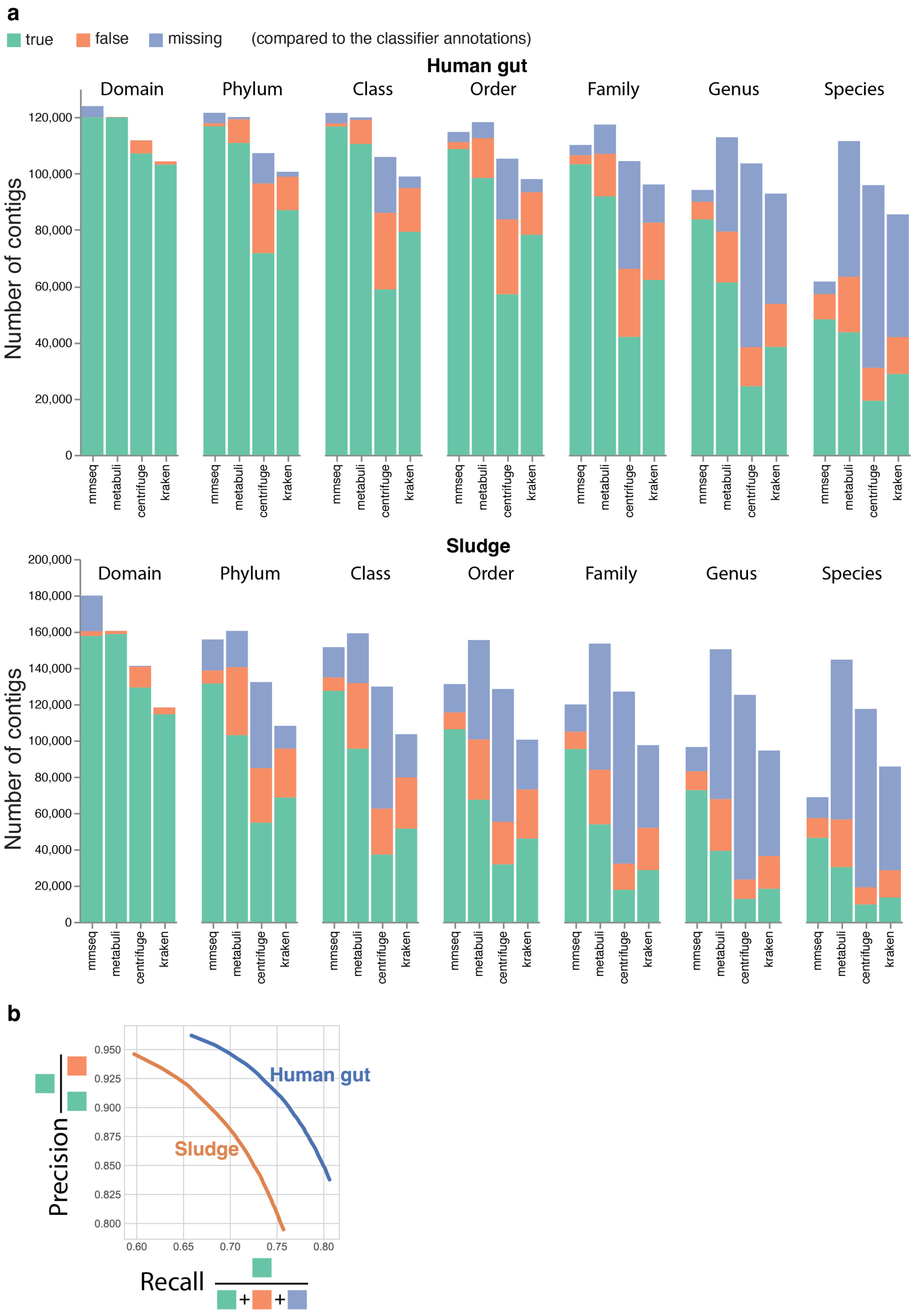


**Supplementary Figure S7 K-fold results for the long-read datasets. a,** The number of true, false and no predictions of Taxometer for four taxonomic classifiers, compared to classifiers annotations, long-read datasets, all domains. **b,** Precision-recall curves for Taxometer scores in the range [0.5, 1] for species labels, long-read datasets.

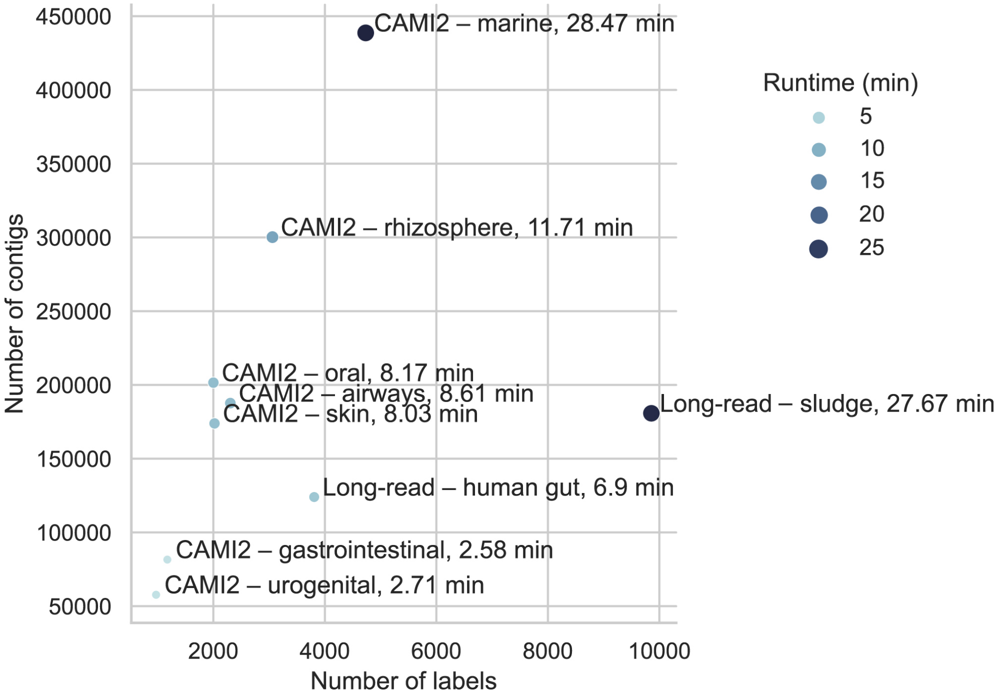


**Supplementary Figure S8 GPU runtimes for all datasets.** X-axis is the number of leaf (species) labels in the taxonomic tree constructed from the classifier annotations for a dataset. Y-axis is the number of contigs in a dataset. The size and hue of a point varies according to the runtime on 1 GPU, in minutes.
